# A novel class of human milk oligosaccharides based on 6’- galactosyllactose and containing *N*-acetylglucosamine branches extended by oligogalactoses

**DOI:** 10.1101/2021.05.27.445961

**Authors:** Franz-Georg Hanisch, Clemens Kunz

## Abstract

Human milk oligosaccharides (HMOs) have attracted much attention in recent years not only as a prebiotic factor, but in particular as an essential component in infant nutrition related to their impact in innate immunity. The backbone structures of complex HMOs generally contain single or repetitive lacto-*N*-biose (type 1) or lactosamine (type 2) units in either linear or branched chains extending from a lactose core. While all known branched structures originate from 3,6-substitution of the lactosyl core galactose, we here describe a new class of HMOs that tentatively branch at terminal galactose of 6’-galactosyllactose. Another novel feature of this class of HMOs was found in linear oligo-galactosyl chains linked to one of the *N*-acetylglucosamine (GlcNAc) branches. The novel structures exhibit general formulas with hexose vs. hexosamine contents of 5/2 to 8/2 and can be designated as high-galactose (HG)-HMOs. In addition, up to three fucosyl residues are linked to the octa- to dodecasaccharides, which were detected in two human milk samples from Lewis blood group defined donors. Structural analyses of methylated glycans and their alditols comprised MALDI mass spectrometry, ESI-(CID)MS and linkage analyses by GC-MS of the derived partially methylated alditol acetates. Enzymatic degradation by application of β1-3,4-specific galactosidase supported the presence of terminal galactose linked β1−6 to one of the two GlcNAc branches.

---

Over the past decade, there is increasing evidence that human milk oligosaccharides (HMO) are of great importance for the health of infants. The beneficial effects of HMOs are discussed to be related to the gut microbiota, inflammatory processes, the immune system or to the development of allergies, and the activity as anti-infectiva ^1–5^. An intriguing aspect is the susceptibility of infants to diseases depending on the type of oligosaccharides they receive. The amounts and profiles of HMOs vary largely during lactation with a total concentration of 5-15 g/L in mature human milk ^6,7^. Today, over 160 structures are elucidated by mass spectrometric and NMR spectroscopic approaches ^8–11^. HMOs are built-up on the lactose core with linear or branched extensions consisting of lacto-*N*-biose and/or lactosamine units which may further be modified by L-fucose and *N*-acetylneuraminic acid. Besides variations in size and branching the most obvious complexity is associated with the Lewis and secretor blood group status of the donor individuals, resulting in Lewis-a, Lewis-b, Lewis-x and Lewis-y structures at the non-reducing termini. Four milk groups can be discriminated based on the Se^+^/se^-^ and Le^+^/le^-^ status: (1) Se^+^/Le^+^ individuals with functional fucosyl transferases FucT2 and FucT3 and Lewis-b positive secretions (Group 1), (2) se^-^/Le^+^ individuals with a defective FucT2 gene (Lewis-a positive; Group 2), (3) Se^+^/le^-^ individuals (blood group H1 positive; Group 3), and (4) the se^-^/le^-^ individuals who lack both functional genes (Group 4) ^12–14^. The different distribution, and hence, the presence or absence of characteristic HMO in the milk of women belonging to the various groups is intensively being discussed to affect various diseases ^2,13,15,16^.

The structural characteristics of classical HMOs is the extension of lactose with GlcNAc at the core galactose and formation of a β1-3 linkage. Branches are introduced at the same sugar by addition of a second GlcNAc linked β1-6. We here describe a completely novel set of branched human milk oligosaccharides, High-Galactose (HG)-HMOs that are tentatively formed on a 6’-galactosyllactose trisaccharide.

## Experimental Section

### Human milk oligosaccharide fractions

Individual human milk samples were provided by healthy women of defined Lewis blood groups and secretor status and processed for enrichment of oligosaccharides in the whey fractions according to previously published protocols ^17^. The fractions under study were obtained during size-exclusion chromatography on BioGel P4. The relevant fractions in the high-mass region of the eluates from a Lewis-b/secretor and a Lewis-a/non-secretor individual were designated as HMO-2 and HMO-107, respectively.

### Reduction of oligosaccharides

Oligosaccharides were treated prior to methylation by reduction with sodium borodeuteride (10 mg/ml 2M aqueous ammonia, 100 µl) for two hours at ambient temperature. After work-up residual salt was removed prior to methylation by solid-phase extraction on carbograph columns (see below).

### Beta-galactosidase digestion of oligosaccharides

Fraction HMO-2 was digested by treatment of 10 µg dried sample with β-galactosidase from bovine testes (10 mU, Prozyme, Invitrogen, Karlsruhe, Germany) in 20 µl of 0.1 M citrate/phosphate buffer, pH 6.0 (Invitrogen) at 37°C overnight. The reaction mixture was desalted by solid-phase extraction on graphitized carbon cartridges.

### Solid-phase extraction on Carbograph cartidges

The samples were desalted and separated from protein/peptide contaminants by solid-phase extraction on graphitized carbon (Extract-Clean columns, Carbograph, 150 mg, 4.0 ml, Alltech, Nettetal, Germany).

### Methylation of glycans

In brief, the methylation of extensively dried samples was performed according to the method described by Anumula and Taylor ^18^. After solubilization in 100 µl dry DMSO (1 vol), methyl iodide was added (0.5 vol) immediately followed by 1 vol of finely powdered sodium hydroxide in dry DMSO. The mixture was incubated for 15 min at ambient temperature before addition of 0.3 ml chloroform and three consecutive extraction steps with 0.2 ml water. The dried chloroform phase was taken up in 100 µl of methanol.

### Profiling of methylated glycans by MALDI mass spectrometry

Methylated glycans in methanol were applied onto a MALDI stainless steel target (Bruker) together with an equal volume of matrix (α-cyano-4-hydroxy-cinnamic acid, saturated solution in 50% acetonitrile/0.1% aqueous TFA). MALDI-MS analysis was performed in the positive ion reflectron mode (HV acceleration 25 kV) on an UltrafleXtreme MALDI-TOF-TOF mass spectrometer (Bruker Daltonics) using the acquisition software FlexControl 3.3 and the data evaluation software FlexAnalysis 3.3 (Bruker Daltonics).

### Partial sequencing of glycans by post-source decay MALDI-MS

MALDI-MS/MS was performed by laser-induced dissociation (LID) in the post-source decay (PSD) mode. Mass annotation was performed in FlexAnalysis and peaklists were generated after manual inspection of the entire mass range for correct isotope peak annotation. The molecular masses of the alditols (M+Na) were searched in standard mass lists, and finally identified by MS/MS using the fragmentation annotation tool provided in the GlycoWorkBench platform.

### Direct inlet ESI-(CID) mass spectrometry of lithium adduct ions

Samples containing 20 µg of methylated oligosaccharide in 200 µl methanol/water (doted with lithium sulfate), 1:1, were directly infused at a flow rate of 3 ml/min via an integrated syringe pump into the Sciex TripleTOF 6600 equipped with a Duo Spray ion source (AB Sciex Germany GmbH, Darmstadt, Germany). Gas flows were set to 10L/min (source gas 1 and 2), and 25 L/min (curtain gas). The source temperature was 50°C, the voltage setting +5500V (declustering potential: 80V). Collision energies were 10V in MS1, and variable between 30V to 60V in MS2. For most experiments the optimum collision energy in MS2 was found at 45V. Spectra were recorded with scan times of 250 msec (MS1), and 100 msec (MS2). The mass range in MS1 was m/z 600-2400, in MS2 from m/z 100-2300.

### Monosaccharide composition and linkage analyses by GC-MS

Oligosaccharides (10 µg) were methanolysed in 1M methanolic HCl for 16 h at 70°C, re-*N*-acetylated and trimethylsilylated as described ^19^. Pyranosidc desoxyhexoses, hexoses and hexosamines were detected by monitoring of the total ion current (TIC) and specific ion traces at m/z 204 and 173.

### Linkage Analysis by GC-MS

Oligosaccharides (10-20 µg) were methylated as described above and converted to the partially methylated alditol acetates (PMAAs) as described previously ^3^. PMAAs were separated and identified by GC-MS on a Fison MD800 equipped with a Rx-5Sil MS capillary column (0.25 ID) using a linear temperature-gradient from 100-280°C (10°C/min). In Addition to the TIC, selected ion traces were monitored at m/z 118 (most hexoses and desoxyhexoses) and at m/z 159 (*N*-acetylhexosamines).

## Results

### Structural characterization of oligosaccharides in fraction HMO-2 and HMO-107 by MALDI-TOF-MS and MS/MS

#### MS1 analysis of HMO-2

Fractions HMO-2 and HMO-107 from Bio-Gel-P4 chromatographies of oligosaccharides isolated from milk of a secretor-(HMO-2) or non-secretor (HMO-107), Lewis-positive individual were initially characterized to contain tetra-to octasaccharides as major components (**Figure 1**). Neither HMO-2 nor HMO-107 contained compounds with relative masses of m/z 527.16 (native) or 681.33 (methylated) that would correspond to trihexoses (not shown). Analysing oligosaccharides in these fractions by HPAE-PAD chromatography revealed further unequivocal evidence for the absence of galactosyllactoses (see Figure S5).

**Figure 1.**
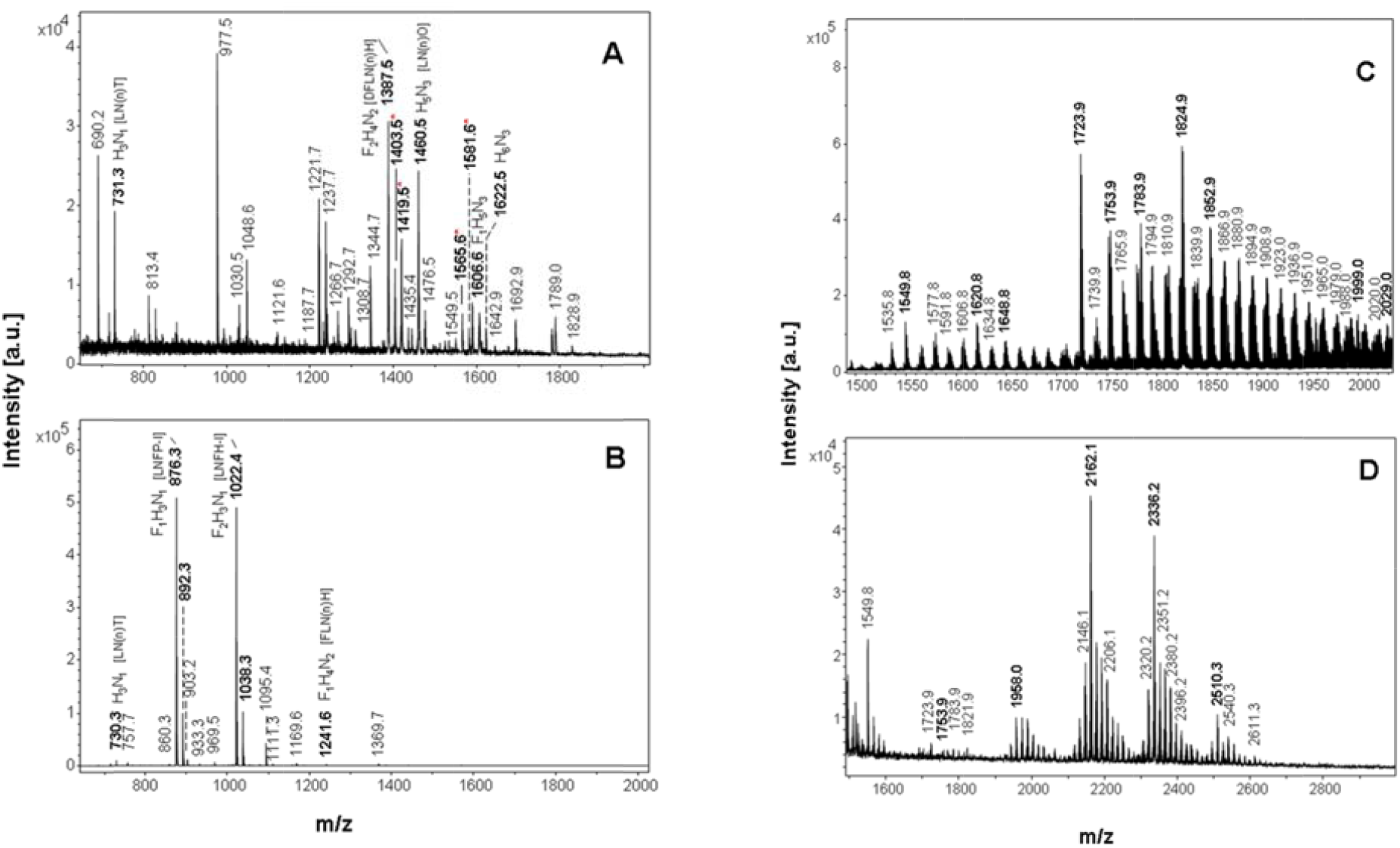
MALDI mass spectrometry survey spectra of native and permethylated oligosaccharides in fractions HMO-2 and HMO-107. Positive ion spectra were recorded in the reflectron mode for native oligosaccharides (A,B) or methylated oligosaccharides in fractions HMO-107 (A,C) or HMO-2 (B,D). Relative masses of signals corresponding to oligosacchari**d**es were highlighted in boldface. Signals corresponding to masses of known HMOs were tentatively annotated in brackets, signals corresponding to novel HMOs were marked by asterisk.

After methylation of fractions HMO-2 and HMO-107 the presence of minor components at low levels became apparent during MALDI mass spectrometry (see **Table 1**). These minor components with masses ranging between m/z 1754-2540 were characterized by high relative proportions of hexoses vs. hexosamines (H5N2 to H8N2) deviating in this respect from the general formulas of known milk oligosaccharides, which can be written as HnNn + 2H. In fraction HMO-107 four species of the new class were detected as native compounds (Table S1) at m/z 1403.5 (F1H5N2), 1419.5 (H6N2), 1565.5 (F1H6N2), and 1581.5 (H7N2), which correspond to the methylated derivatives with molecular masses M+Na at m/z 1753.9 and 1783.9, respectively (only trace signals were detectable at m/z 1958.0 and 1988.0). All other species were detected in fraction HMO-2 after permethylation, but were undetectable in MALDI mass spectrometry as native compounds.

**Table 1.**
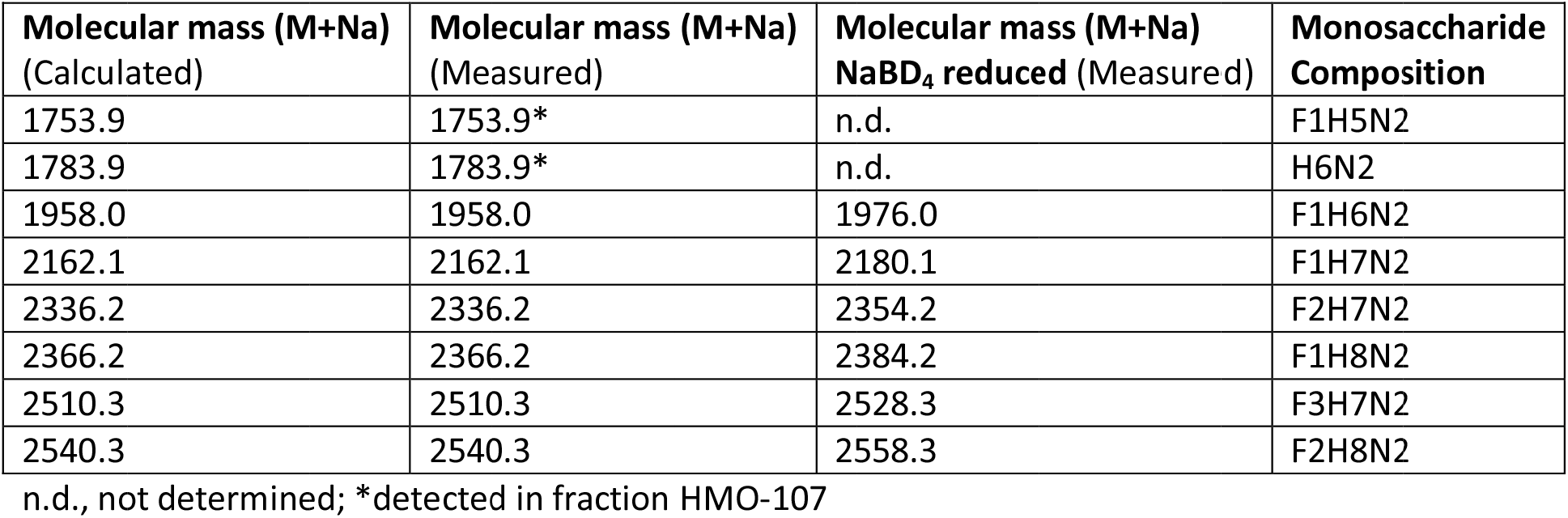
Novel HG-HMO species detected in human milk oligosaccharide fractions HMO-2 and HMO-107 by MALDI mass spectrometry.

The precursor ion detected in fraction H**M**O-107 at m/z **1754** (M+Na) revealed characteristic fragmentation patterns in PSD-MALDI-MS (**Figure 2**) that were indicativ**e** of a branched octasaccharide (F1H5N2).

**Figure 2A.**
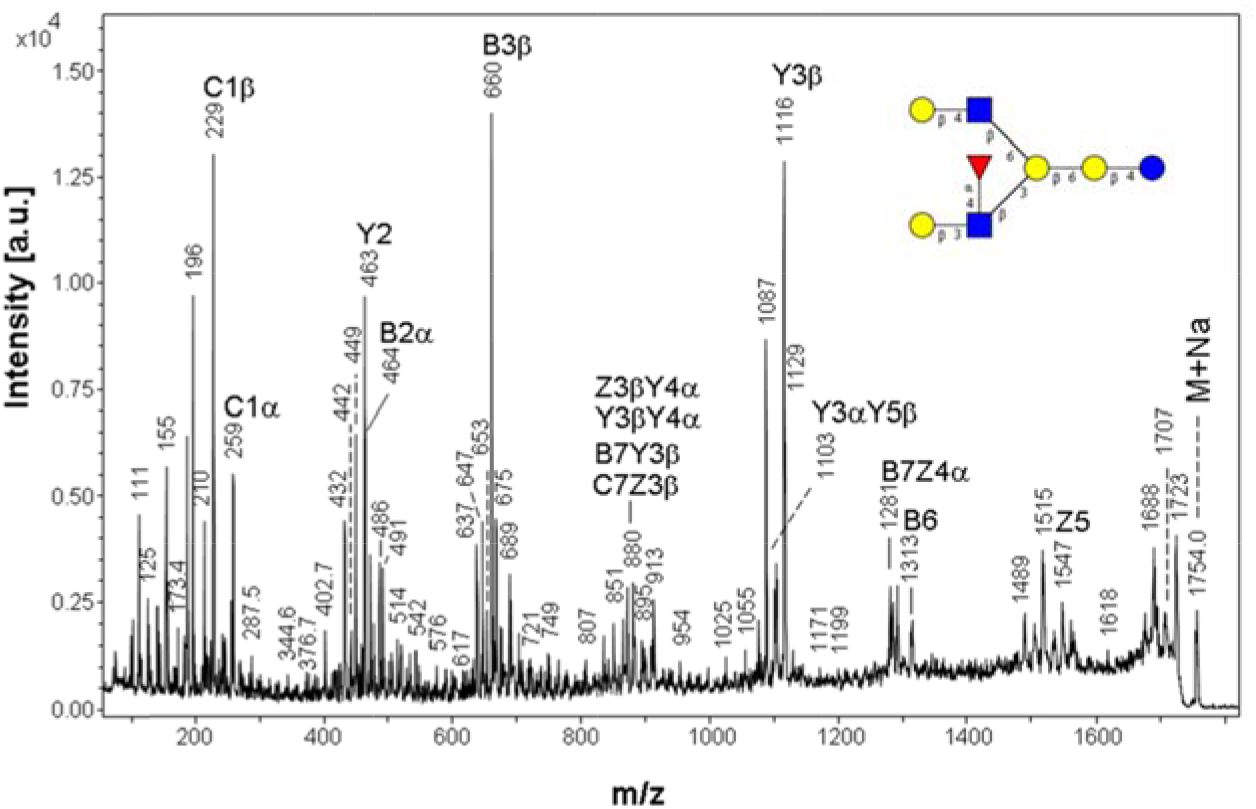
Figure 2 MALDI-MS/MS by post-source decay analysis of a octasaccharide (m/z 1754) detected in fraction HMO-107. Postive ion MALDI-MS of permethylated fraction HMO-107 revealed a parent ion mass (M+Na) at m/z 1754 that was selected for PSD analysis. Annotation of fragment ions was performed according to predictions calculated by the GlycoWorkBenc**h** tool for fragment ions of the B,C,Y, and Z series. Designation of fragment ions followed the nomenclature of Domon and Costello based on the tentativ**e**ly assigned 6-Gal/3-Gal branch pattern. Only a selection of ion masses and their annotations are shown in the graph together with a tentative structural model that integrates also data from methylation/linkage analysis. A d**i**scrimination between 6-Gal and 3-Gal branches was not possible on the basis of MALDI-MS/MS fragmentation.

Fragments from the non-reducing terminus were detected at m/z 229 (C1), and 660 (B3β) indicating terminal Hex-(dHex-)HexNAc or Gal-(Fuc-)GlcNAc, and at m/**z** 464/486 (B2α), indicating a second terminal unit Hex-HexNAc (Gal-GlcNAc). Two ions **a**rise from double fragmentation with formation of Z3Z3 at m/z 617 and Y3Y3 at m/z 653, b**o**th indicating the presence of a trihexosyl unit at the reducing terminus. The ion at m/z 463 could correspond to a Y2 fragment, and would support the trihexosyl unit. Although it may als**o** arise from fragmentation of a Hex-Hex-HexNAc br**a**nch at 6Gal of a lactose-core, it could not yield the double fragmentation ions at m/z 617 and 653. The ouble fragmentation ion at m/z 880 is expectedly formed also from the alternative structural model. Further evidence for a Hex-HexNAc branch and a trihexosyl core unit comes from the ions at m/z 1103 (Y3αY5β), 1116 (Y3β), and 1291 (Y3α). A unique fragment, which can unequivocally be assigned to a branched B6 ion is detected at m/z 1313. The ion at m/z 1547 should correspond to Z5β and confirms again the trihexosyl core unit.

To overcome some of the limitations associated with the MALDI-MS/MS approach and to get deeper **i**nto structural details of the novel HMOs with respect to substitution sites and branching configuration, we chose an alt**e**rnative mass spectrometric approach genera ing cross-ring and internal fragmentation of lithium adduct ions under high-en**e**rgy collision dissociation conditions. While protonated oligosaccharides produce mainly glycosidic cleavages, the respective metal ion adducts yield increased cross-ring cleavage even under conditions of low energy CID (**Ref**). Lithium adduct ions in fraction HMO-107 contained singly and doubly charged species at m/z 1738 and 872, respectively, that **c**orrespond to the sodium adduct species at m/z 1754. MS2 of these ions revealed besides ions of the B, C, Y, and Z series intense and partially unique cross-ring fragments (see ions marked by an asterisk in **Figure 2B**).

**Figure 2B.**
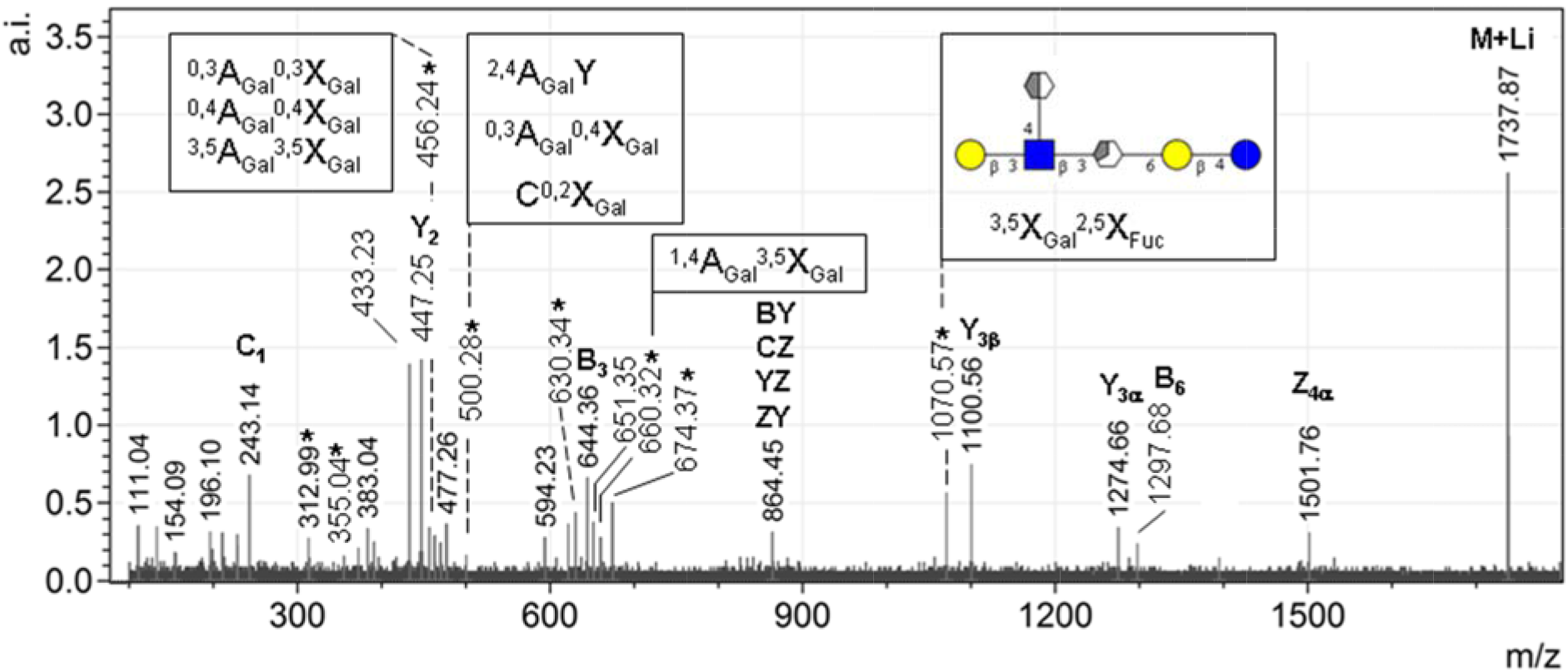
Direct inlet ESI-(CID)MS2 of lith um adduct ions of methylated octasaccharide m/z 1738. Beside fragments of the B, C, Y, and Z series cross-ring fragments were annotated. Unique cross**-**ring fragments revealing important structural information with respect to branching configuration were highlighted by insertion of structural interpretations. Ambiguous cross-ring fragments were either marked by an asterisk or by an asterisk in brackets (isobaric with double cleavage ions).

While most of these cross-ring fragments are ambiguous with r spect to their structur l assignments, several species were unique and allowed unequivocal structu**r**al identification (see ions annotated with cross-ring fragment specifications). The series at m/z 456, 500, 660, 674 and 1070 is in support of the branch configuration shown in Figure 2, which locates the type 1 chain to the 3-linked branch. Refe**r**ring to the most intense and unique fragment at m/z 1070, it corresponds to ^3,5^X_Gal_ ^2,5^X_Fuc_, a hexasaccharide fragments, which h**a**s lost the 6-linked branch (see insert in Fig. 2B).

The precursor ion detected in fraction H**M**O-2 with a relative molecular m**a**ss (M+Na) at m/z 1958 was of minor abundance and not further characterized by MS/MS. The most prominent signals in the high-mass range of survey spectra were those at m/z **2162**, and **2336** (the signal at m/z 2510 and 2528 was of minor abundance, but partial sequence data could be obtained by MS/MS of the deutero-reduced compound, refer to **Figure S1**).

The parent ion at m/z 2162 corresponding to a decasaccharide (F1H7N2) revealed again several ion series which support a branched backbone structure with two non-reducing termini: m/z 433 (C2) and 660 (B3) indicating a Fuc-Gal-GlcNAc branch and m/z 463 (C2) and 912 (C4) indicating a Gal-Gal-Gal-GlcNAc branch (**Figure 3A**: refer to structure S1 representing ions shown in re).

**Figure 3.**
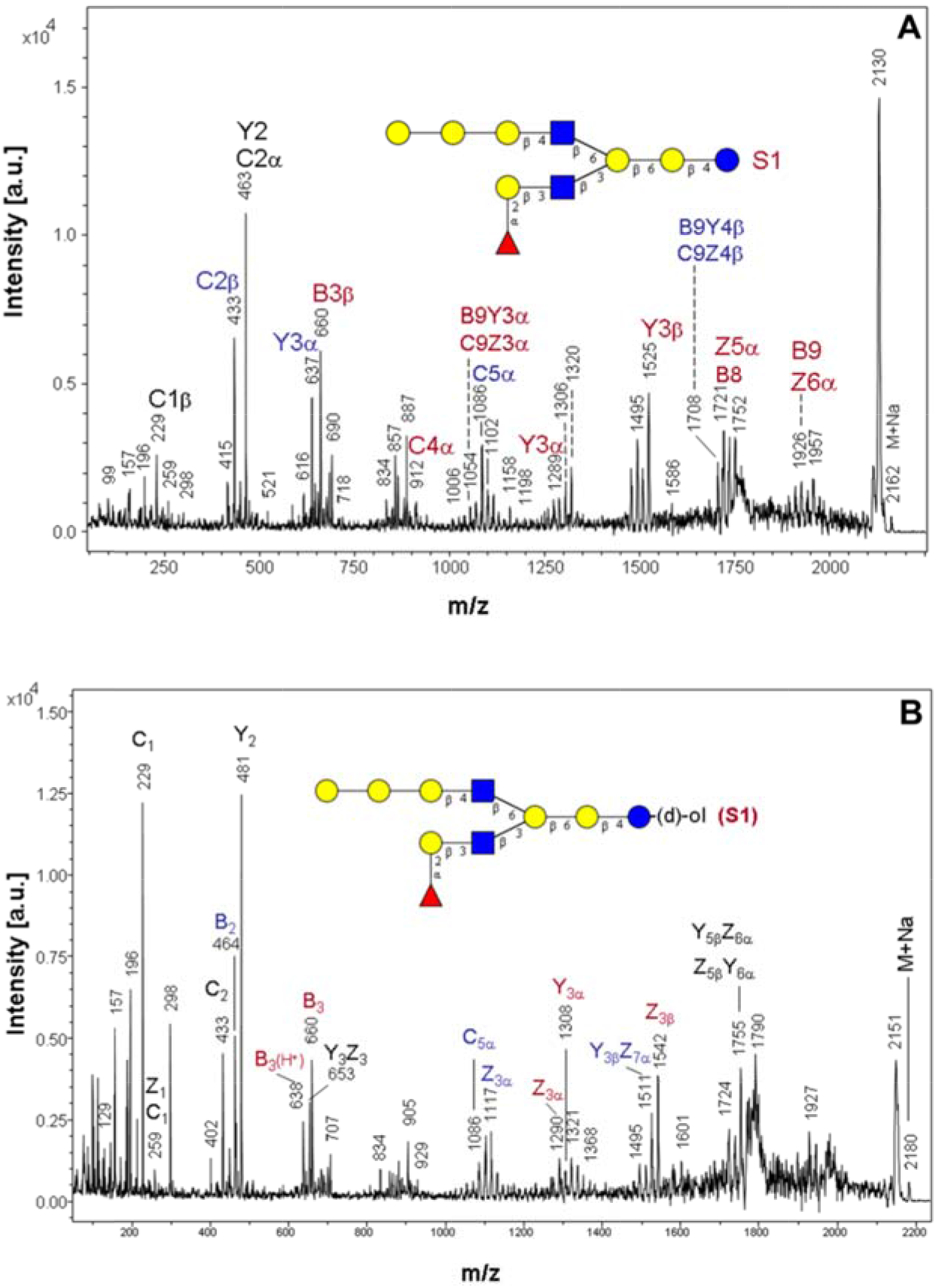
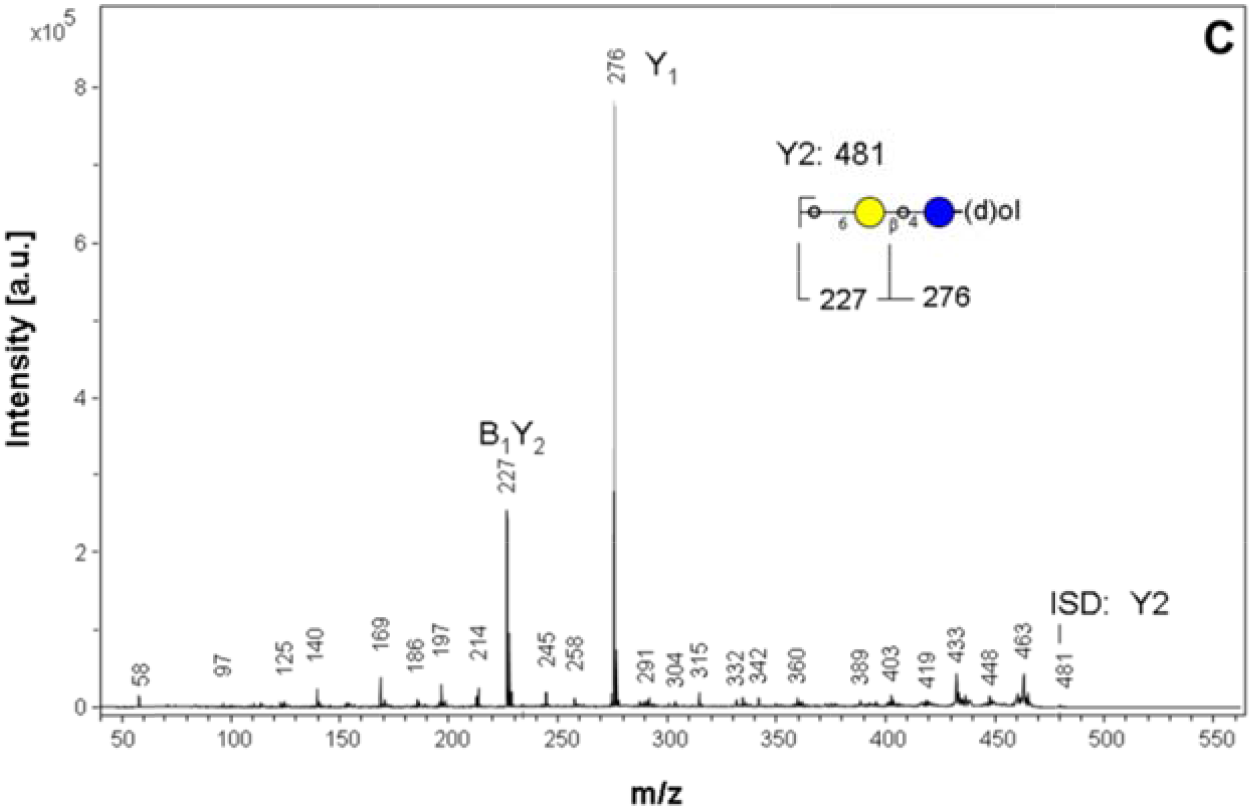
MALDI-MS/MS by post-source decay analysis of a decasaccharide (m/z 2162) and its deutero-reduced alditol (m/z 2180) detected in fraction HMO-2. Postive ion MALDI-MS of permethylated fraction HMO-2 revealed a parent ion mass M+Na) at m/z 2162 that was selected for PSD analysis (A). After eutero-reduction of the fraction the parent ion mass shifted to 2180 (B) and the respective fragmentation spectrum allowed a discrimination of oligohexosyl fragments from the reducing or non-reducing ends of theoligosaccharide. A discrimination between 6-Gal and 3-Gal branches was not possible on the basis of ALDI-MS/MS fragmentation. The PSD-fragment at m/z 481 detected after deutero-reduction and representing the trihexosyl core is also formed by In-Source Decay and was selected for PSD fragmentation analysis (C). Isomeric structure model S2 (D). Fragment annotation (see Figure 2).

Ions supporting the existence of a trihexose at the reducing end were registered at m/z 1054 corresponding either to B9Y3α or to C9Z3α (assignments to 6-Gal/3-Gal branches are only tentative). While several fragment ions are ambiguous with respect to the structural assignment, the ions at m/z 1524 and 1752 can unequivocally be identified as Y3β and Z4β ions, respectively, which strongly support the trihexosyl core (see also the B9/Z9α ion at m/z 1925). Finally, the unique fragment ion at m/z 1956 can be assigned as Z5**β** ion, which is in support of the structural model shown in Figure 3 (structure S1).

The strongest argument for the novel structural features of components in fraction HMO-2 comes from MALDI-MS/MS of deutero-reduced methylated HMO-2 oligosaccharide alditols, as these reveal a series of unique fragments, which cannot arise from lacto**s**e-based branched structures. After borodeuteride reduction of oligosaccharides in fraction HMO-2 the parent ion corresponding to m/z 216 was registered at m/z 2180 (**Figure 3B**). While most fragments containing the non-reducing regions of the oligosaccharide were identical to those registered for the glucoaldose compound (see above), a unique series of Y and Z type fragments were indicative of the respective deuterated gl**u**citol compound. In particular, th**e** ions at m/z 481 (Y2), 1290 (Z3α), 1308 (Y3α), and 1542 (Z3β) strongly support the existence of a trihexosyl core. On PSD fragmentation of the in-source fragment at m/z 481 two major ions at m/z 227 (B1Y2) and 276 (Y1) were formed (**Figure 3C**).

Few ions correspond to an isomeric compound, which differs from structure S1 by the position of fucose (**Figure 3A**: refer to relative masses shown i blue; stru**c**ture S2). The ions at m/z 637 (C3), 1068 (B5α), and 1086 (C5α) indicate that an isomeric decasaccharide carries fucose at the oligogalactosyl branch (**Figure 3D**: isomeric structure model S2).

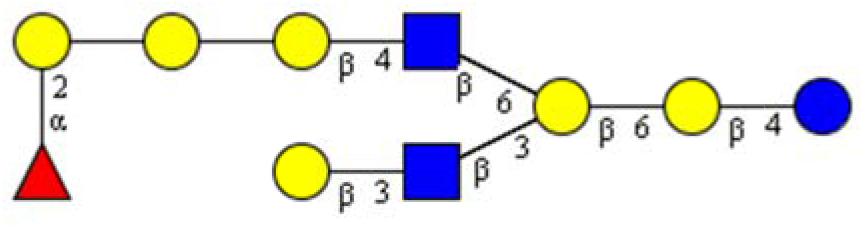

The parent ions at m/z 2336 and its corresponding deutero-reduced derivat**i**ve registered at m/z 2354 revealed similar pat erns of fragment ions supporting again two fucose subs itution isomers represented by PSD fragments masses shown in red (st ucture S1) or in blue (structure S2) in **Figure 4**.

**Figure 4.**
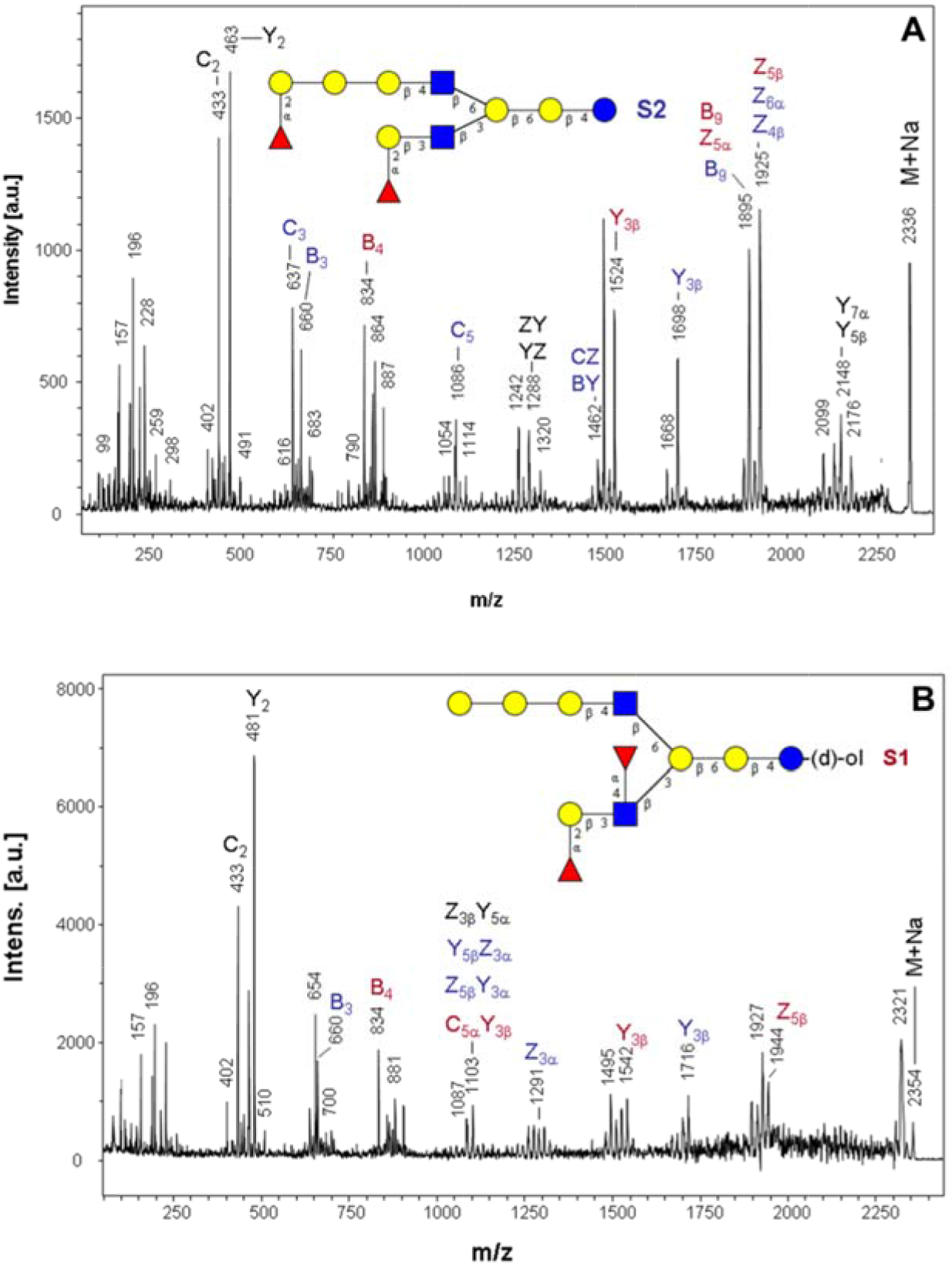
MALDI-MS/MS by post-source decay analysis of an undecasaccharide (m/z 2336) and its deutero-reduced alditol (m/z 2354) detected in fraction HMO-2. Postive ion MALDI-MS of permethylated fraction HMO-2 revealed a parent ion mass (M+Na) at m/z 2336 that was selected for PSD analysis (A). After deutero-reduction of the fraction the parent ion mass shifted to 2354 (B) and the respective fragmentation spectrum allowed a discrimination of oligohexosyl fragments from the reducing or non-reducing ends of the oligosaccharide. Fragment annotation (see Figure 2). A discrimination between 6-Gal and 3-Gal branches was not possible on the basis of MALDI-MS/MS fragmentation.

Ion series at m/z 834 (B4), and 1524 (Y3β) indicate the presence of structure S1, in which two fucosyl residues substitute the type 1 lacto-*N*-biose at the 3-Gal branch (Lewis-b epitope), whereas ion series at m/z 637 (C3), 660 (B3), 1086 (C5), and 1698 (Y3β) can be assigned to the structure S2 model, in which the two fucosyl residues are linked to terminal galactoses in both branches (**Figure 4A**). PSD-fragmentation of the deutero-reduced derivative revealed identical series of relative masses for the non-reducing terminus and fragments with a mass shift of +18 for those arising from the reducing terminus (see for example the alternative Y3β ions in **Figure 4B**). The existence of a trihexosyl core is again supported by the strong Y2 ion at m/z 481.

### Monosaccharide and methylation/linkage analyses of fractions HMO-2 and HMO-107: detection of 6-Gal glycan nodes

A monosaccharide composition analysis by GC-MS of trimethylsilylated 1-*O*-methylglycosides revealed fucose, galactose, glucose, and *N*-acetylglucosamine as components in the molar ratios 0.5/2.4/1.0/1.4 (HMO-107) or 2.1/2.4/1.0/2.9 (HMO-2), respectively (normalization was done relative to Glc as 1.0)(refer to **Figure S2**). Fucose is present in HMO-107 at lower relative abundancies compared to fraction HMO-2. Galactose was found at higher relative molar abundancies Gal/Glc than 2.0 in both fractions. A linkage analysis of oligosaccharides was performed for both fractions (HMO-2 and HMO-107) after partial enrichment on graphitized carbon (**Figure 5**).

**Figure 5.**
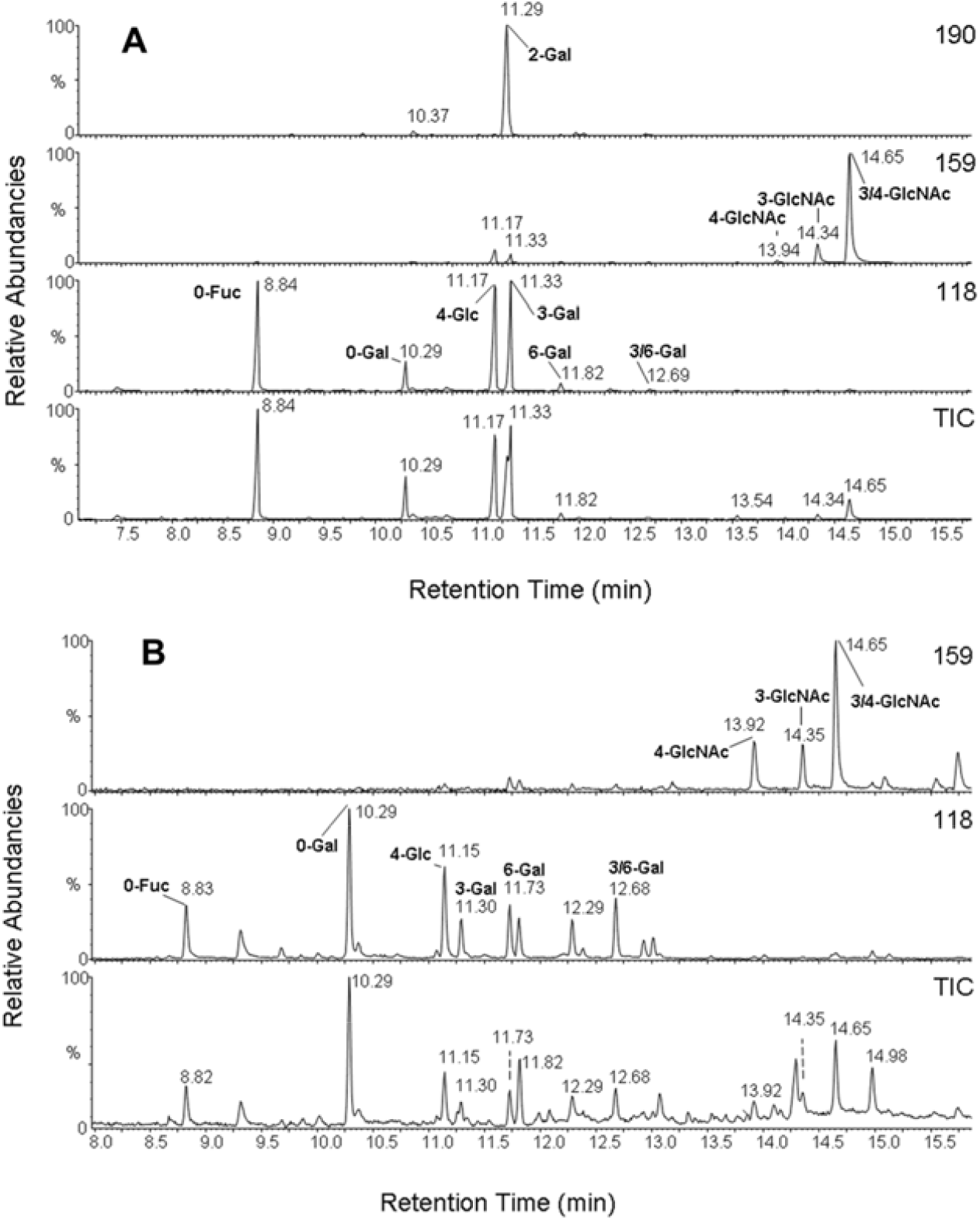
Linkage analysis by GCMS of partially methylated alditol ace ates derived from methylated fractions. (A) PMAA from fraction HMO-2 were separated by gas-chromatography and the Total Ion Current (TIC) or selected ion traces formed by EI mass spectrometry were monitore at m/z 118 (desoxyhexoses, hexoses), at m/z 159 (*N*-acetylhexosamines) or at m/z 190 (2-substituted hexoses). (B) PMAA from fraction HMO-107 were separated and monitored as above. Refer to selected EI mass spectra and GCMS profiles of a PMAA standard mixture in **Figure S3**.

Fraction HMO-2 was still dominated by LNFP-I and LNDFH-I as major c**o**mponents. Accordingly, only those partially methylated alditol acetates (PMAA) that are structurally unrelated to these major components can be regarded as arising from the minor components corresponding to the novel H -HMOs (**Figure 5A**). These latter PMAAs comprize 1,3,5,6-tetra-*O*-acetylated galactose (,6-Gal) indicating a 3,6-branch agalactose, and a 1,5,6-tri-*O*-acetylated galactose indicatin 6-Gal, which could be found within the non-reducing oligo-galactose chains or correspond to the lactosyl-galactose. The detection of 1,4,5-tri-*O*-acetyl-N-acetylglucosaminitol (4-Gl NAc) is indicative of a type 2 lactosamine chain. In fraction HMO-107 the proportion of novel HG-HMO species was significantly higher (20-30%) according to the MALDI-MS survey spectra of native oligosaccharides. Linkag analysis revealed the presence of PMAA representing 0-Fuc, and 0-Gal as structural elements (glycan nodes) of the non-reducing termini (refer to the signals at 8.9 and 10.3 min, respectively, in **Figure 5B**). A series of PMAAs or glycan nodes ^20^ can be expected to be derived from major HMOs corresponding to general structural features (HnNn+2H): 4-Glc, 3-Gal, 3,6-Gal, 4-GlcNAc, 3-GlcNAc, and 3,4-GlcNAc. A high proportion of terminal galactose (0-Gal) together with significant amounts of 3,6-Gal (refer to the signal at 12.7 min) support the presence of branched oligosaccharides with non-reducing type 1 (lact-*N*-biose) or type 2 (lactosamine) chains, which can be fucosylated at subterminal glucosamine residues (refer to **Figure S3**). One PMAA (1,5,6-tri-*O*-acetylated galactose) is not generally expected to occur in HMOs of higher molecular masses, as it indicates the presence of 6-Gal (**Figure S3**). This glycan node is confined to the trisaccharide levels, as it was found exclusively in galactosyl lactoses (6’Gal-Lac)(Yamashita and Kobata, 1974).

### Enzymatic degradation of HMO-2 oligosaccharides with β-galactosidase

To reveal information on the linkage types of galactoses in the oligo-galactose branch of HG-HMOs fraction HMO-2 was digested with the β3,4-specific galactosidase from bovine testes. The control substrate lacto-*N*-tetraose (m/z 926) was quantitatively digested to the corresponding trisaccharide GlcNAc1-3Gal1-4Glc detected at m/z 722 (**Figure S3**). On digestion of oligosaccharides in fraction HMO-2 the HG-HMO series did not show any significant conversion to degalactosylated species, indicating that terminal galactose should not be linked 1-3/4 to the subterminal sugar (see oligosaccharide masses in boldface in **Table 2**).

**Table 2.**
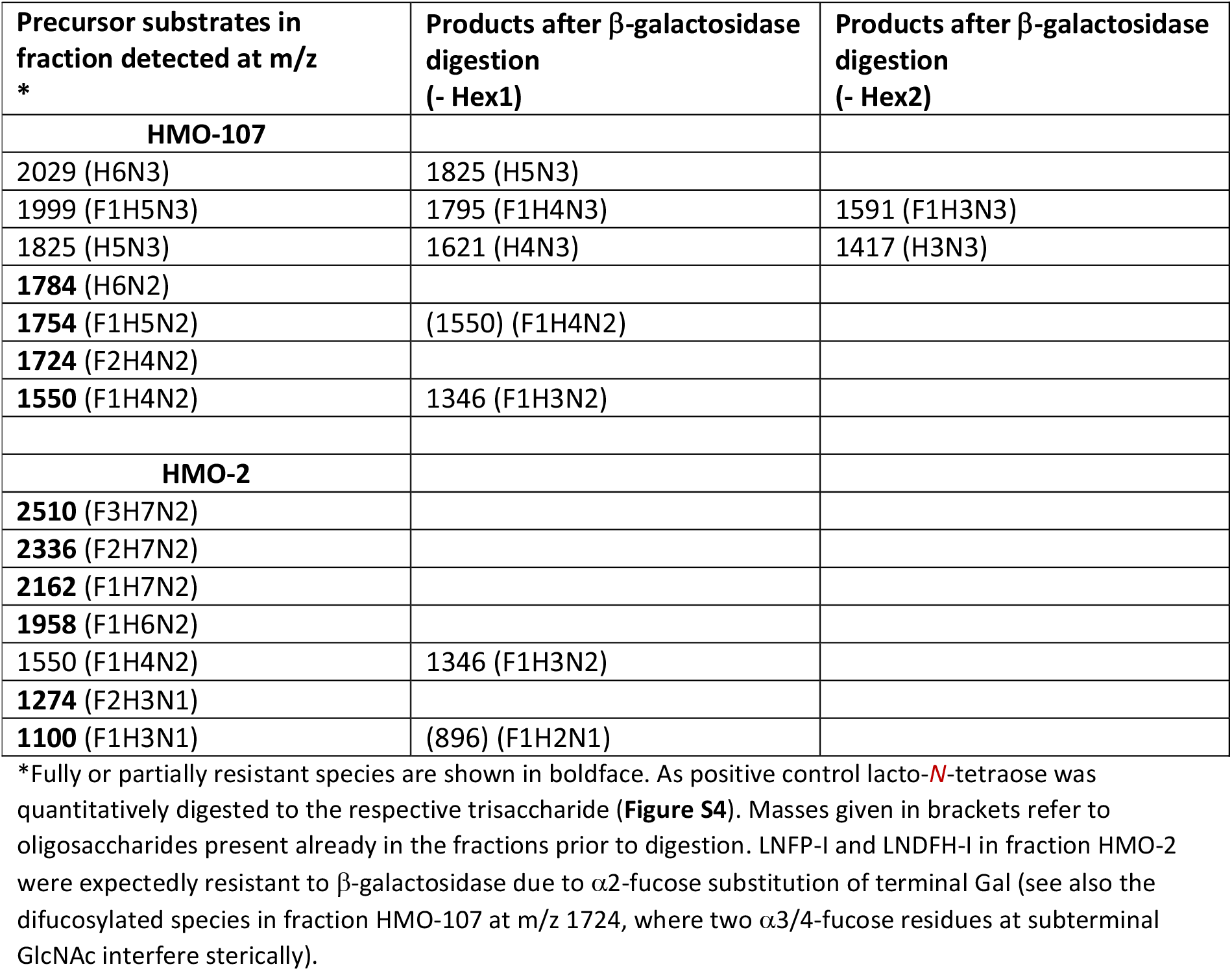
MALDI mass spectrometry of oligosaccharides in fractions HMO-2 and HMO-107 prior to and after β-galactosidase digestion.

On digestion of oligosaccharides in fraction HMO-107 several species were quantitatively converted to –Hex1 or -Hex2 products (**Table 2**), whereas others (masses in boldface) were largely or fully resistant to enzymatic degradation. This holds true for HG-HMO compounds at m/z 1784 and 1754. The mass detected after digestion at m/z 1550 (a potential –Hex product of the compound at m/z 1754) should be identical to an isomer detectable prior to digestion. The compound at m/z 1724 corresponds to a β-galactosidase-resistant oligosaccharide (F2H4N2) with one fucose in each GlcNAc branch. The major signal at m/z 1824 corresponding to an octasaccharide with “classical” backbone structure is fully degraded to the –Hex1 and –Hex2 products.

## Discussion

On screening high-mass HMO fractions from various mothers for their binding capacity to noroviral capsids ^3^ we observed unexpected molecular masses and obscure fragmentation patterns in some samples when analysed by MALDI-MS (**Table S1**). A thorough investigation led to the postulation of a new class of HMOs, which we propose to call “HG-HMO”. The “classical” HMOs are characterized by the general formula H_n_N_n_+2H, with only glucose (in the lactose core) and galactose and *N*-acetylglucosamine as constituents of repetitive linear or branched lacto-*N*-biose or lactosamine units, which may be modified by the attachment of one or more fucose and/or NeuAc residues. The new class of oligosaccharides in human milk exhibits a distinct general formula: H_n+m_ N_n_+3H. The structures are based on a trihexosyl core followed by two *N*-acetylglucosamine branches of which one carries a number of m further hexoses corresponding to oligo-galactoses. Those repeating Gal moieties in one branch are not present in classical HMO. A further striking feature is also that both branches may be fucosylated with up to three fucose moieties. So far, similar structures have not been reported neither to be present in milk of any other species.

Although the discovery of HMO goes back more than six decades to the pioneering work of Richard Kuhn, Jean Montreuil and others ^9,21^ it is surprising that this new HMO class has not been discovered earlier. One reason is the low amount in the lower milligram to upper microgram range per liter affording enrichment and analysis with advanced mass spectrometric methods. A further reason for their escape from detection may be their low desorption efficiencies in MALD ionization as native compounds (see, for example, the undetectability of HG-HMOs in fraction HMO-2 prior to methylation). Only by applying MALDI-TOF-MS and MS-MS of methylated oligosaccharides a conspicuous fragmentation could be seen (**Figures 1-4**). Further detailed investigations comprised methylation/linkage analysis by GC-MS in combination with specific exoglycosidase digestions to reveal insight into terminal structural elements. As we revealed structural information for only two individual milk samples, very few generalisations can be made. All structures of the HG-HMOs are characterized by a trihexosyl core identical with galactosyllactose (Gal-Lac), most likely 6’Gal-Lac. The presence of a 6’Gal-Lac core is supported by methylation linkage analyses, but also by the concentrations of the potential biosynthetic precursor in milk samples, in particular from non-secretor individuals ^22^. A further common structural feature is the presence of oligo-galactosyl chains extending from one of the GlcNAc residues. The patterns of HG-HMOs in the two fractions under study differ in some aspects that refer to the size (larger structures were seen in the secretor sample HMO-2) and number and positions of fucosyl residues (higher degree of fucosylation in the secretor sample, localization at 2-Gal in HMO-2, whereas HMO-107 reveals substitution of fucose at 4-GlcNAc according to methylation analysis).

Comparing classical HMO and HG-HMO with oligosaccharides in milk of various animals it is evident that only human milk contains such a high number of diverse HMOs. Although each animal species has its own milk oligosaccharide pattern, the number of components is usually rather low and quantitative data for most components are not available. For more information on milk oligosaccharides in various species and their differences we refer to the very thorough publications by Urashima and coworker ^23–25^.

Occasionally, there are reports of the presence of β-galactosyl-lactoses (Gal1-6Gal1-4Glc, Gal1-4Gal1-4Glc, and Gal1-3Gal1-4Glc) in human milk ^11,22,26^. However, in humans these trisaccharides, only in minute amounts detectable and rarely quantifiable, were never found to be extended, whereas in marsupial milk extended unbranched oligo-galactosyl-lactose chains were found, besides unusual branches at lactosyl-cores, which were detected in the “novo”-oligosaccharide series, where the 6-Gal branch starts with a galactose ^23,24^. Distinct from the new HG-HMO, however, is that these components, e. g. in the Tammar Wallabys neither contains the GlcNAc branching nor the fucosylation at either branching arm. To differentiate the new HG-HMO from the commercially used GOS it is important to underline that GOS contains a variable number of galactose moieties as the only monosaccharide in their galacto-chains ^27,28^. Hence, GOS do not have any similarity neither to classical HMOs nor to HG-HMOs

Functions of the new HG-HMO are not known yet for obvious reasons, but the hitherto unknown structural peculiarities increase further the already high potential of HMO for very specific effects shown in many preclinical studies. The complexity of oligosaccharides in human milk might be one of the reasons why formula-fed infants are more prone to diseases than breast-fed infants ^29^.

To summarize, we identified a new class of HMO called High Galactose-HMO characterized tentatively by (i) a 6’-galactosyllactose trisaccharide, (ii) a GlcNAc branching at 3,6-Gal, (iii) a variable number of repeating Gal moieties in one of the branches and (iv) a variable fucosylation pattern. These novel structures have not been reported to be present in milk of any animal species nor in any biological material.

## Abbreviations

ESI-(CID)MS: electrospray-(collision-induced dissociation) mass spectrometry
GC-MS: gas chromatography-mass spectrometry
HMO: human milk oligosaccharide
HPAE-PAD: High pH Anion Exchange chromatography with Pulsed Amperometric Detection
MALDI-MS: matrix-assisted laser desorption ionization mass spectrometry

## Acknowledgements

The current project was supported by a grant of the Köln-Fortune programme (to FGH). Authors acknowledge the kind technical assistance of Cordula Becker preparing milk samples for studies with classical HMOs. We are thankful to Dr. Stefan Müller from the Center for Molecular Medicine Cologne, who kindly supported us with the performance of direct-inlet ESI-(CID)MS.

## Supporting information

*Table of contents*

Title page

### Legends to supplementary figures and tables

Figure S1: MALDI-MS/MS by post-source decay analysis of a deutero-reduced dodecasaccharide alditol (m/z 2528) detected in fraction HMO-2

Figure S2: Monosaccharide composition analyses by GCMS of oligosaccharides in fractions HMO-2 and HMO-107

Figure S3: EI mass spectra of PMAA derived from permethylated oligosaccharides in fraction HMO-107

Figure S4: MALDI-MS1 spectra measured after β-galactosidase digestion of oligosaccharides in fractions HMO-107 or HMO-2

Figure S5: HPAE-PAD chromatography of fraction HMO-2 (A) and HMO-107 (B)

Table S1: Native and methylated glycans of the H_n+m_ N_n_+3H type detected in human milk oligosaccharide fractions HMO-2, -7, -107, -127, -142

## For Table Of Contents Graphics

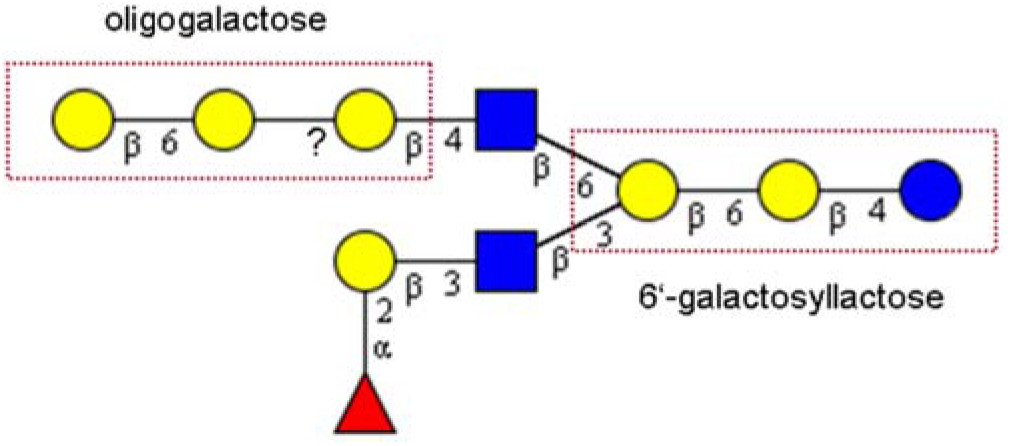

